# Learning to read transforms phonological into phonographic representations: Evidence from a Mismatch Negativity study

**DOI:** 10.1101/2024.09.10.611672

**Authors:** Chotiga Pattamadilok, Shuai Wang, Deidre Bolger, Anne-Sophie Dubarry

**Affiliations:** Aix Marseille Univ, CNRS, LPL, Aix-en-Provence, France; Institute of Language, Communication and the Brain (ILCB); Aix Marseille Univ, CNRS, CRPN, Marseille, France

## Abstract

Learning to read changes the nature of speech representation. One possible change consists in transforming phonological representations into phonographic ones. However, evidence for such transformation remains surprisingly scarce. Here, we used a novel word learning paradigm to address this issue. During a learning phase, participants were exposed to unknown words in both spoken and written forms. Following this phase, the impact of spelling knowledge on spoken input perception was assessed at two time points through an unattended oddball paradigm, while the Mismatch Negativity component was measured by high density EEG. Immediately after the learning phase, no influence of spelling knowledge on the perception of the spoken input was found. Interestingly, one week later, this influence emerged, making similar sounding words with different spellings more distinct than similar sounding words that also share the same spelling. Our finding provides novel neurophysiological evidence of an integration of phonological and orthographic representations that occurs once newly acquired knowledge has been consolidated. These novel phonographic representations may characterize how known words are stored in literates’ mental lexicon.

## Introduction

The ability to process spoken language is considered to be universal. This intuitive assumption implies that literate and illiterate individuals share at least the most fundamental speech processing skills that allow them to recognize spoken words. In line with this view, *all* existing psycholinguistic models of spoken word recognition have been built upon data that were exclusively collected in literate populations. In other words, any potential differences in how literate and illiterate listeners recognize spoken words and in the nature of representations stored in their mental lexicons have been disregarded (see ^1,2^ for general discussions on the impact of literacy acquisition on human language and cognition).

The present study investigated the nature of the representations stored in the phonological lexicon of literate listeners. It addressed a specific question of whether these representations are purely phonological. The question emerged from the existing literature which suggests that the impact of learning to read is far beyond a simple connection between the two language codes. More specifically, this connection, which is promptly established after being exposed to the spoken and written input, could induce more profound and long-term changes within the spoken language system itself. For instance, it has repeatedly been reported that spoken words that had inconsistent spellings (e.g., /aɪt/ can be spelled ‘ite’, ‘ight’, or ‘yte’) took more time to be recognized than spoken words that had consistent spellings e.g., /ʌɡ/ is always spelled ‘ug’)^3–6^. This increase in reaction time was associated with an increase in amplitude of a negative-going ERP waveform around 320-350 msec after stimulus presentation; a time-window that coincided with the onset of lexical access^7,8^. Interestingly, further examination of the cortical source of this ERP signal showed that the so-called “orthographic effect” was located in the left supramarginal gyrus (SMG), a classic phonological area. No evidence for activation in the left-ventral occipital cortex (vOT, also known as the Visual Word Form Area) coding orthographic information was found^7^. The same conclusion was obtained in a study that used transcranial magnetic stimulation to interfere with the orthographic consistency effect by interrupting the function of either the left-SMG or the left-VOT^9^. The result showed that the reaction time advantage of orthographically consistent over inconsistent words was removed when the stimulation was delivered to the left SMG whereas it was maintained when delivered to the left-vOT.

Another type of evidence for a modulation of brain activity within spoken language areas following reading acquisition comes from comparisons of brain activity in participants with different levels of reading expertise. Using this approach, Dehaene and colleagues^10^ compared brain activity (measured by fMRI) of literate, ex-illiterate and illiterate Brazilian/Portuguese participants during spoken word recognition and sentence listening. The authors found that the activity within the planum temporale increased gradually with participants’ reading skill. However, it is worth noting that Hervais-Adelman et al.^11^ who addressed a similar issue in Hindi speakers who were either illiterate or literate in a non-alphabetic script did not observe the same result. These diverging results further suggest that the impact of literacy on the modulation of brain activity should not be interpreted as an all-or-none phenomenon. In addition to inter-individual variability such as reading expertise, the characteristics of the writing system itself might also play an important role.

Despite the various origins of variability, the reported modulations of brain activity within the spoken language system following reading acquisition seems to fit with the general idea that phonological knowledge is progressively restructured through language experience^12,13^. As discussed by Muneaux and Ziegler ^14,15^, although the lexical restructuring framework mainly focuses on the role of vocabulary size, it is plausible that reading acquisition, which is one of the most critical language experiences, also contributes to restructuring the spoken language system to some extent^16–18^. One form of lexical restructuring is the emergence of abstract *phonographic* representations. These abstract representations reflecting an integration of phonological and orthographic features of language input have been mentioned or alluded to in the literature on several occasions ^14,19–22^. Yet, direct evidence of such transformation remains surprisingly scarce. One noteworthy finding was reported by Bakker et al.^23^ who conducted behavioral experiments to investigate the formation of novel word memories across modalities. In their experiments, participants learned either the spoken or written version of novel words. After the learning phase, within- and cross-modal integration of the novel words to the mental lexical was behaviorally assessed by measuring the degree to which visually or auditorily acquired novel words engaged in a lexical competition with existing phonological or orthographic neighbors. The authors found that the novel words learned in the spoken form needed 24h before entering into phonological and orthographic neighbor competitions. Novel words learned in the written form needed 24h before entering into orthographic neighbor competition but one week before entering into phonological neighbor competition. This behavioral evidence of a spontaneous generation of cross-modal representations from language input that had been learned in a specific modality indicates that, once an individual had become literate, the information contained in his/her mental lexicon might become multimodal. Interestingly, this claim is compatible with neurophysiological evidence reported by Pattamadilok et al.^24^. In this electroencephalography (EEG) study, the authors examined the nature of the “phonological” lexical representations in literate participants by recording auditory Mismatch Negativity (MMN) responses in an unattentive oddball paradigm where participants were exposed to sequences of standard and deviant spoken words. Standard and deviant stimuli always ended with the same phonological rimes. Critically, some deviant stimuli also shared the same rime spelling with the standard stimuli while others did not (e.g., standard: /tRi/, spelled “tri”; deviant with the same rime spelling: /kRi/, spelled “cri”, deviant with different rime spelling: /pRi/, spelled “prix”). Although the manipulation of orthographic congruency remained implicit in this auditory oddball protocol, the amplitude of the MMN was found to be modulated by participants’ knowledge of the spoken words’ spelling: A higher MMN amplitude was observed in the situation where the rime spellings of the standard and the deviant spoken words were mismatched than when they were matched. Since MMN is an index of automatic detection of changes in both concrete and abstract features of the sensory input^25–27^, its sensitivity to the orthographic feature of spoken input suggests that this feature might be integrated into the memory traces of spoken words.

However, the stimuli used in Pattamadilok et al.^24^ were known spoken words with known spellings, and thus, did not allow the authors to reveal any transformation of the nature of the underlying representations at different learning stages. In other words, even if there was an integration of phonological and orthographic features, it would have already taken place well before the experiment. The present study aimed to fill this gap by conducting a two-phase study. In the *novel word learning phase*, participants learned the spoken and written forms of unknown words. Contrary to past research, this protocol allowed us to control two critical factors, i.e., the participants’ knowledge about pronunciation and spelling of the stimuli and the precise moment at which this knowledge was presented to the participants. In the second phase, the nature of the mental representations associated with the newly acquired words was examined through an *unattended oddball paradigm*. In this phase, participants were passively exposed to the spoken version of the novel words that they had learned, and their brain activity was recorded with high density EEG. As in the previous study^24^, our critical manipulation was the orthographic congruity of the learned words presented as standard and deviant in the oddball paradigm (e.g., standard: /izo/, spelled “izôt”; deviant with the same rime spelling: /ivo/, spelled “ivôt”, deviant with different rime spelling: /iʒo/, spelled “ijaux”). The difference between the auditory MMNs obtained in the orthographically incongruent and orthographically congruent situation was examined at two time points, i.e., immediately after the learning phase and one week later^23^, without additional learning between the two experimental sessions. The second EEG recording session was added to the protocol following the predictions made by the Complementary Learning Systems model of memory consolidation^28–30^. According to this framework, lexical acquisition operates in two stages. The first stage corresponds to a rapid formation of episodic memory. The episodic representations are stored in the hippocampal system as independent, context-specific, and non-integrated sparse codes. This initial learning stage was followed by a slower neocortical learning during which novel information is gradually integrated into existing memory networks by forming connections between the novel words and orthographically, phonologically, or semantically related existing words. Hence, the resulting consolidated representations are assumed to be encoded in widely distributed networks and are no longer modality specific^23^. A number of studies provided evidence for the role of overnight sleep in the transfer of newly acquired knowledge stored as episodic representations into lexicalized long-term memory, with the duration of the consolidation period varying from a few hours to several days^23,31–34^. So far, this framework has mainly been used to explain how new knowledge is integrated into the existing body of knowledge ^32,35–37^. Here, we argued that the same integration process might also apply to the combining of information from different input modalities associated with the new knowledge.

The present study aims at revealing the nature of the underlying representations associated with the newly learned spoken words, and how these representations evolved across the two learning phases. We hypothesized that if the memory traces of the novel spoken words only contained the phonological features, the MMN would not be sensitive to the orthographic congruency between the standard and the deviant stimuli. If, in contrast, the orthographic features were integrated into these memory traces, as reported in Pattamadilok et al.^24^, MMN amplitude should increase in the situation where the standard and deviant spoken stimuli were orthographically incongruent. Importantly, the evolution of the MMN across the two time points should provide insightful information about the dynamic of the underlying representations. Specifically, if the impact of spelling knowledge resulted from an activation of the episodic memories of word spellings acquired during the learning phase, it should be detected during the first EEG session which took place immediately after the learning phase, when the episodic traces were still recent. However, if the impact of spelling knowledge resulted from an integration of the phonological and orthographic features that occurred during the lexicalization process, it should be detected during the second EEG session, once the newly acquired knowledge had been consolidated and lexicalized.

## Results

### Behavioral data

The letter detection task applied during the learning phase showed that, after the initial exposure and the first dictation task, all participants had successfully learned the spellings of all novel words. The averaged accuracy score obtained in the task was 96.3% (range: 87.5% - 98.7%). All participants obtained 100% correct responses in the subsequent dictation tasks conducted on Day 1 and Day 8 after the EEG recording session, which suggested that the spelling knowledge was correctly learned and remained accurate one week after the exposure to the novel words.

### Electroencephalographic data

#### ERP analysis

For each experimental session and each participant, peak amplitude, Area Under Curve (AUC) and peak latency of the MMNs obtained for the orthographically congruent and incongruent conditions were extracted from three electrode clusters (anterior central, central and posterior central) selected based on extant literature. Prior to analyzing the effects of orthographic congruency and experimental session, we assessed whether the grand average MMN (which included all experimental conditions and sessions) was observed as expected in each electrode cluster. The comparison of the MMN peak amplitudes extracted from the time-window of interest with the peak amplitude extracted from the baseline period showed significant differences in the central [t(127) = -7.63, p < .0001] and posterior central clusters [t(127) = -5.03, p < .0001] but not in the anterior central cluster [t(127) = 1.76, p > .08]. Based on this observation, only the central and posterior central clusters were retained for further analyses.

Following the above preliminary analysis, we examined the effects of orthographic congruency and session. The analyses conducted on MMN peak amplitude and AUC showed the same pattern of results. The ANOVA summary of the LME showed a significant interaction between orthographic congruency and session for both dependent variables [F(1, 224) = 4.47, p = .036 and F(1, 224) = 4.47, p = .036 for peak amplitude and AUC, respectively]. No other main effects or interactions showed any significance (p values > .094). Based on the significant interaction, we specifically tested our main hypothesis on the influence of orthographic knowledge on spoken language perception by comparing the data obtained in the orthographic congruent and orthographic incongruent conditions within each experimental session (i.e. Day 1 and Day 8). As illustrated in Figure 1, in the first experimental session, MMNs obtained in the orthographically congruent and incongruent conditions were statistically equivalent [t(231) = 0.64, p > .51, 95% CI of the difference between the two MMNs -0.34, 0.17 and t(231) = 0.80, p > .42, 95% CI -6.35, 2.69 for peak amplitude and AUC, respectively]. Interestingly, when the MMNs were measured one week later, we found that MMN amplitude induced by the orthographically incongruent deviant was larger than that induced by the orthographically congruent deviant [t(231) = -2.30, p = .022, 95% CI -0.55, -0.042 and t(231) = -2.14, p = .033, 95% CI -9.43, -0.40 for peak amplitude and AUC, respectively]. The difference between the pattern of results obtained in the first and the second experimental session was caused by a significant increase of MMN amplitude generated by the orthographically incongruent deviant measured one week after the learning session [t(231) = -2.64, p = .009, 95% CI -0.60 - 0.087 and t(231) = -2.54, p = .012, 95% CI -10.33, -1.30 for peak amplitude and AUC, respectively]. No difference between the MMNs generated by the orthographically congruent deviant was found when the two sessions were compared [t(231) = 0.30, p > .76, 95% CI -0.29, 0.22 and t(231) = 0.40, p > .68, 95% CI -5.44, 3.59 for peak amplitude and AUC, respectively].

**Figure 1.**
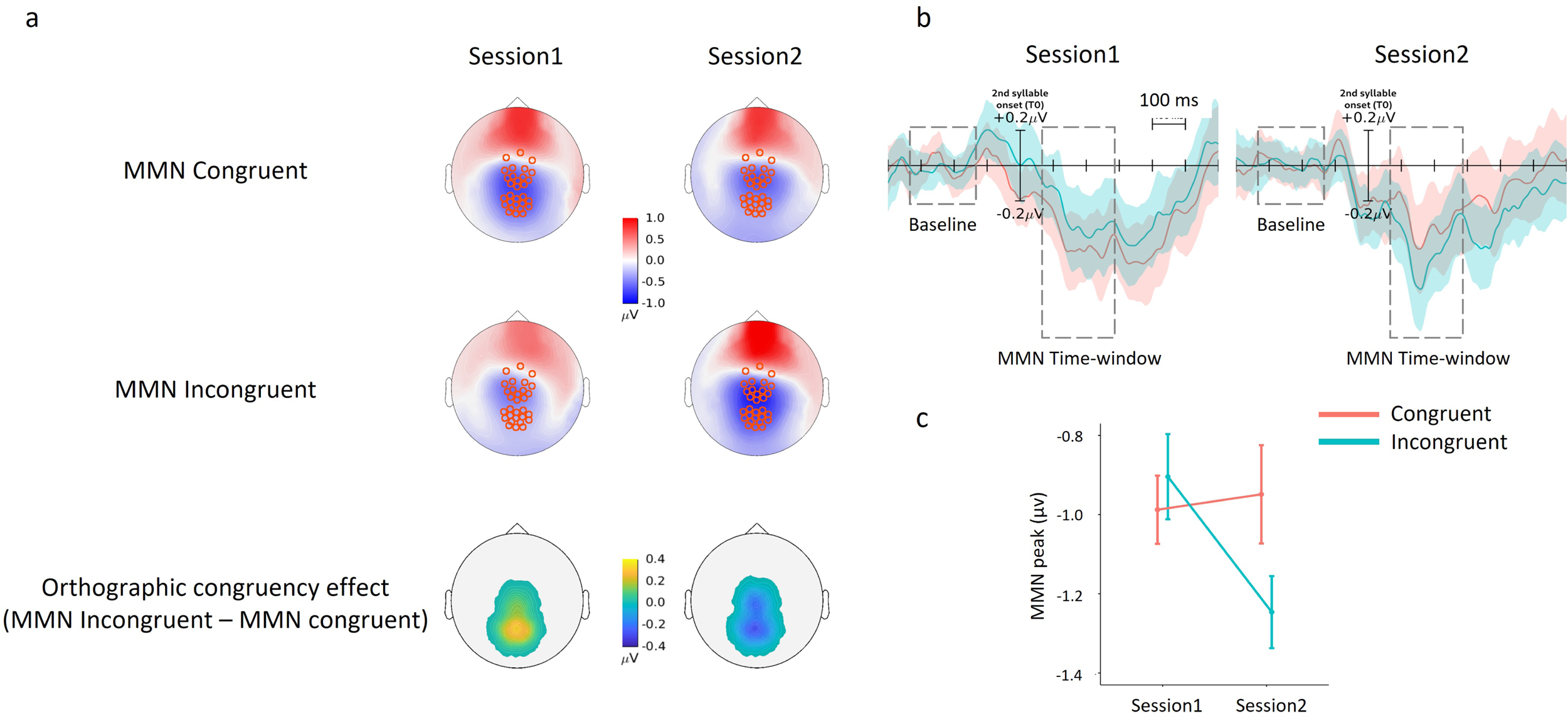
MMN topographic maps, waveforms, and peak amplitude. **a**. Full scalp topographies of the MMNs obtained in the orthographically congruent and incongruent conditions in session 1 and session 2 (the electrodes from the central and posterior-central clusters highlighted in red circles). Topographical view of the difference between the two conditions at the central and posterior central clusters showing the impact of orthographic congruency on the MMN amplitudes. **b**. Averaged ERP waveforms (T0 = second syllable) extracted from the central and posterior central clusters, for each condition in each session. **c**. MMN peak amplitudes extracted from the central and posterior-central electrode clusters, for each condition in each session. Error ribbons in B and error bars in C represent standard error of the means.

Finally, the analysis conducted on MMN latency only revealed a significant main effect of cluster [F(1, 224) = 5.30, p = .022]. The latency of the MMN recorded in the central cluster was slightly shorter than that observed in the posterior central cluster (139±20 ms vs. 145±26 ms). No other main effect or interaction presented a significant result (p values > .13).

#### Source localization analyses

To explore the generators of the orthographic congruency effect observed in the ERP analysis, we compared the EEG reconstructed sources in the incongruent condition compared to the congruent condition recorded during the second experimental session. The statistical analysis of the full cortex revealed stronger neural responses in the incongruent condition, located in the left middle temporal gyrus (MTG), left middle frontal gyrus (MFG) and left superior parietal lobule (SPL), whereas a decreased neural response in the incongruent condition was observed in the left fusiform (FUS) (p values < 0.001 unc. with cluster size > 10; see Figure 2 and Table 1S in supplementary material).

**Figure 2.**
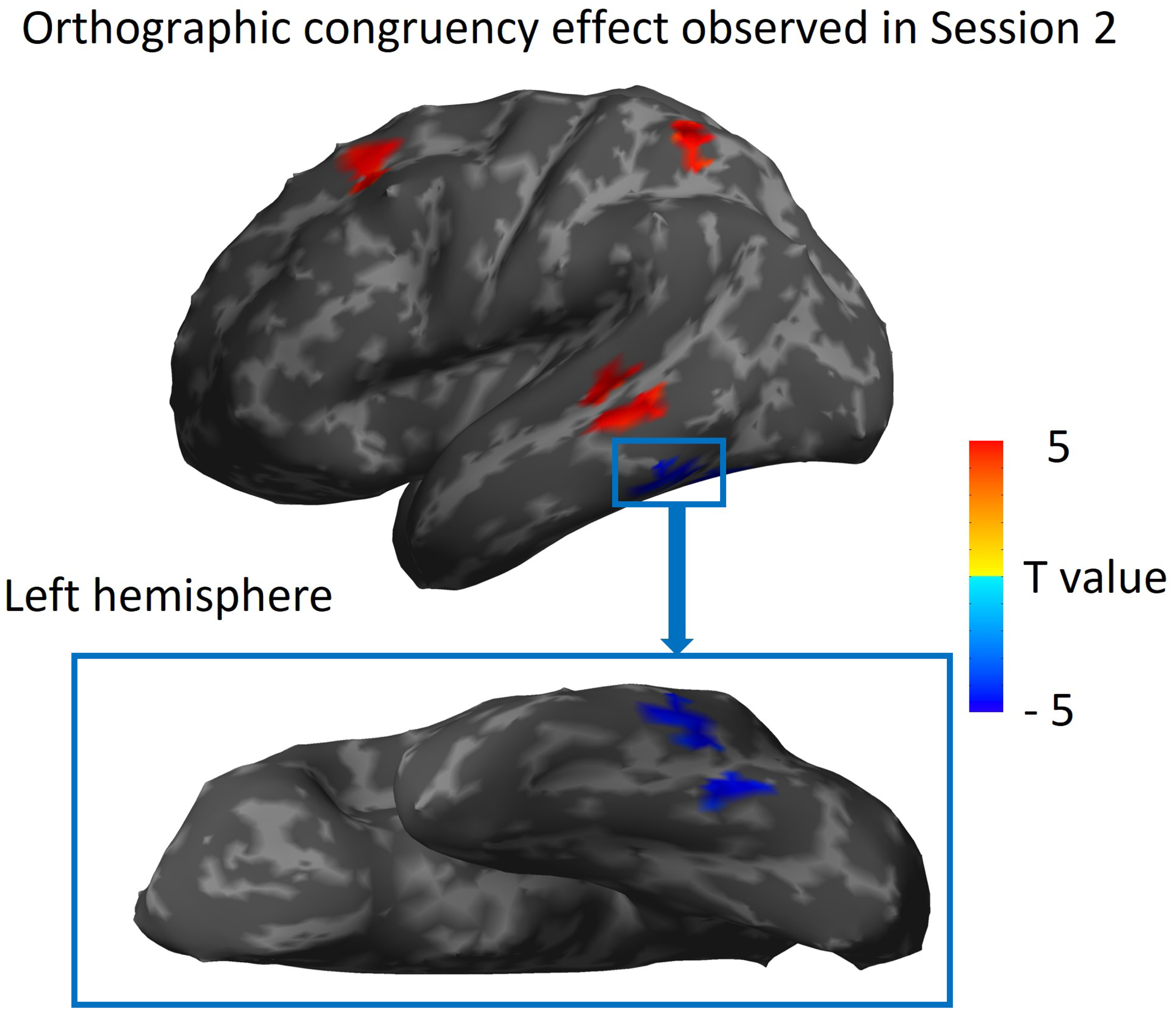
Group level statistical source map of the orthographic congruency effect (incongruent - congruent) observed in the second experimental session. The computation was conducted in the MMN time-window. Left middle temporal cortex, left middle frontal gyrus and left superior parietal lobule showed stronger neural responses in the incongruent compared to the congruent condition while the left fusiform showed the opposite pattern of result. P values < 0.001 unc. with cluster size > 10.

**Table 1:**
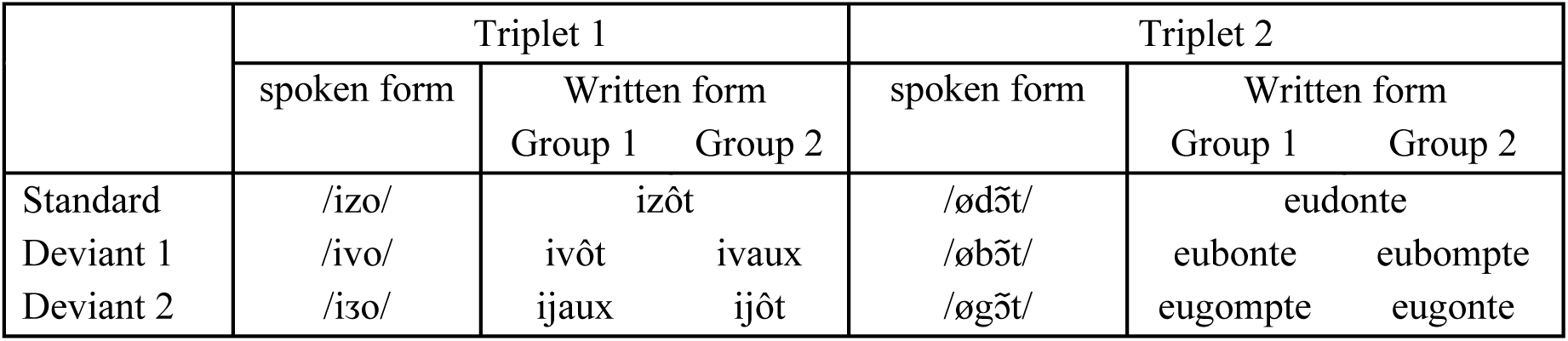
Two novel word triplets and their associated spellings presented to the two groups of participants.

## Discussion

Phonological lexicon is the core element of spoken word recognition models. Yet, despite a significant amount of work devoted to understanding the nature of the representations stored in the mental lexicon, no consensus has been reached regarding the format and the level of abstractness of the representations. The absence of such a consensus could be due, in part, to the plasticity of the speech recognition system itself which, under some circumstances, enables newly acquired knowledge or language experience to modulate the nature of the existing representations and interactions between them (see ^38,39^ for extensive reviews on this topic). Based on the current literature on the impact of reading acquisition on speech processing, the present study further explored the consequences of this plasticity by testing the hypothesis that, once individuals learn to read, the nature of spoken words’ representations might no longer reflect the phonological properties of spoken words alone but would also be contaminated by how spoken words are spelled.

The novel word learning protocol adopted here allowed us to manipulate the phonological and orthographic representations associated with new linguistic inputs, as well as the moment at which these inputs were learned by the participants. After the learning phase, the nature of the novel words’ representations was assessed in a low-level and strategy-free processing context by examining the modulation of MMN recorded in an unattentive oddball paradigm where only the spoken form of the words was presented. The analyses that we conducted aimed to address two questions: First, whether the MMN elicited by the spoken form of the new words was also sensitive to implicit knowledge of the words’ spellings as previously reported on known words^24^. Second, if such sensitivity exists, would it emerge immediately after being exposed to the linguistic inputs or only after a consolidation period.

The analysis of the MMN recorded immediately after the learning phase showed no sensitivity to the manipulation of word spelling even though the participants had correctly acquired and memorized this information. Interestingly, the sensitivity to spoken word’ spelling spontaneously emerged one week later without additional exposure to the stimuli between the two sessions. This sensitivity was reflected by a significant increase of the MMN amplitude (indexed by peak and AUC) in the condition where the standard and the deviant spoken words were orthographically incongruent compared to when they were orthographically congruent. Given that this difference was not observed in the first session with the same stimuli, it could not be attributed to potential uncontrolled differences in the acoustic features of the stimuli used in the different experimental conditions. Further exploration of the cortical sources of the orthographic congruency effect obtained in the second session showed a stronger brain activity in distributed brain areas including the left MTG, left MFG, and left SPL. Interestingly, we also observed a reduction of brain activity in the left FUS.

What does this change in the sensitivity to spoken word spelling across the two experimental sessions tell us about the nature of the representations stored in the mental lexicon? As mentioned in the Introduction, our finding is considered within the Complementary Learning Systems (CLS) framework^28–30^. The behavioral data collected in both letter detection and dictation tasks clearly indicated that all participants correctly learned the spoken and written forms of the novel words after a short learning period. According to the CLS, at this initial learning stage, these two input modalities would be stored as separate entities. Our finding is coherent with this claim: The MMN observed immediately after the learning phase was sensitive only to the phonological aspect of the spoken inputs, thus suggesting that the episodic memory traces of the spoken words presented during the unattentive oddball task remained modality specific at the initial stage. Most importantly, the pattern of MMN response recorded one week later revealed a change in the nature of the representations. Although our protocol did not allow us to identify the precise moment when the change occurred, it clearly suggests that the modality-specific episodic memory traces of the spoken word forms (and most probably of the written word forms) had been consolidated and transferred to long-term memory within the interval of one week. The consolidation process led to an integration of the phonological and orthographic features of the inputs, thus, rendering the underlying representations of spoken words multimodal (or amodal, see^23^) as revealed by the MMN response. This observation is coherent with both the MMN pattern reported in Pattamadilok et al.’s study^24^ conducted on known words and a behavioral study reported by Bakker and colleagues^23^ who argued that “*…in the process of consolidation, novel words acquire an amodal lemma representation which no longer contains detailed episodic information about initial exposure conditions. This amodal representation would be able to engage in competition both in spoken and printed word recognition, regardless of modality consistency between training and test phases*.” (p. 127). Although no cross-modal lexical competition was assessed in the present protocol, the fact that the MMN elicited by spoken words became sensitive to their spellings is coherent with the presumed transformation of modality-specific word forms to multimodal or amodal representations. The claim regarding converging representations of words that have been learned in the spoken and written modality is also coherent with the conclusion proposed by Partanen and colleagues^40^ who argued in favor of similar cortical dynamics that are involved in lexicalization of novel word forms learned in auditory and visual modalities. Regarding the underlying cortical sources activity related to the consolidation of new linguistic knowledge, Bakker-Marshall and colleagues^41^ reported in an EEG study a modulation of theta oscillations, which was argued to reflect a consolidation process, occurring in the left posterior middle temporal gyrus (pMTG), a region known to be involved in lexical storage^42–45^. While EEG-based source localization does not allow us to draw a strong conclusion on the cortical sources of the effect of interest, the fact that our analysis revealed the left-MTG as one of the regions that showed an increase of activity in the orthographically incongruent condition in the second session is coherent with Bakker-Marshall et al.’s finding. The two other brain regions that showed an increase of activity, i.e., the left-MFG and the left-SPL, are generally considered as parts of the domain-general multiple demand network that supports high-level cognition^46–48^. Their recruitment has also been reported in various linguistic tasks, along with other areas of the phonological network such as the left inferior frontal gyrus and the left temporoparietal region^49–51^. For the time being, one could speculate that the effect observed in these regions could be linked to additional cognitive resources allocated to processing incoming inputs that differed from the existing memory traces in both phonological and orthographic dimensions. Interestingly, in addition to the increase of activity in the regions mentioned above, we also observed a decrease of activity in the left-FUS at the approximate location of the Visual Word Form Area^52^. This activation pattern, which is opposite to the direction of the effect captured by the MMN component, argued against the possibility that the orthographic knowledge might be activated during the task. On the contrary, it seems to reflect a cross-modal inhibition of the visuo-orthographic system during spoken word perception^53^. The observed decrease of left-FUS activity in the orthographically incongruent compared to congruent condition could result from a stronger top-down inhibition from the spoken language system to the visuo-orthographic system in the presence of conflicting information. While awaiting confirmation from future research, the findings from this exploratory source localization point to a consolidation process that recruits brain areas in distributed networks within and outside the spoken language system.

## Conclusion

So far, the potential impact of reading acquisition on speech processing has mainly been explained by the online activation of orthographic representations that occurs while speech is being processed^52,54^. While the evidence for the online activation of orthographic representations is particularly robust in high-level speech processing tasks (see ^55^ for a review), the findings obtained in speech processing tasks that involve lower perceptual stage are less consistent^56–58^, thus leading to a debate about whether the impact of learning to read on speech processing truly exists or simply reflects experimental or task-dependent artifacts. Here, we avoid this pitfall by investigating the impact on orthographic knowledge in a low-level speech processing context where spoken inputs were processed in the absence of focussed attention. Our finding clearly indicates that the impact of reading acquisition on speech processing does not necessarily reflect an online mapping or interaction between the two language codes. In fact, orthographic knowledge has already exerted its influence on speech representations well before participants performed the task, i.e., by rendering speech representations multimodal. This transformation occurs spontaneously outside individuals’ awareness or strategic control. Hence, this new evidence raises the possibility that, once we learn to read, the representations stored in our “phonological” lexical may no longer be purely phonological.

## Methods

### Participants

Sixteen right-handed native French speakers aged 18–32 years (mean 23.3 years) participated as paid volunteers. They were randomly separated into two groups of four men and four women. All had normal hearing and no history of neurological or language disorders. The experiment was performed in accordance with the Declaration of Helsinki and was approved by the local ethics committee (Sud Méditerranée I: RCB 2015-A00845-44). Informed consent was obtained from all participants.

### Stimuli

The novel words to be learned by the participants corresponded to two triplets of disyllabic pseudowords having V_1_-C_2_V_2_ and V_1_-C_2_V_2_C_2_ phonological structures. Within each triplet, one novel word was used as standard stimulus and two as deviant stimuli in the auditory oddball paradigm. The novel words within each triplet differed at the position of the onset of the second syllable (C_2_). Otherwise, they always shared the first syllable (V_1_) and ended with the same phonological rime (V_2_ or V_2_C_2_).

To manipulate the orthographic congruency between the standard and the deviant stimuli, for each triplet, the standard novel word was associated with one spelling while each deviant novel word was associated with two spellings. One of the two spellings was congruent with the spelling of the standard stimulus while the other one was incongruent. As shown in Table 1, for participants in Group 1, the rime spelling of Deviant 1 was orthographically congruent with that of the standard novel word, while the rime spelling of Deviant 2 was orthographically incongruent. The associations were reversed for the participants in Group 2. Overall, the standard stimulus remained constant and each deviant stimulus was associated with both congruent and incongruent spellings across the two groups of participants. This manipulation allowed us to ensure that any differences that might be observed between the MMNs obtained in the orthographically congruent and incongruent conditions were not due to the physical properties of the deviant stimuli (e.g., amplitude, duration, intensity and spectral characteristics).

The spoken pseudowords were each recorded several times using a female native French speaker voice in a soundproof room. All recorded stimuli were digitized at a sampling rate of 44.1 kHz with 16-bit analog-to-digital conversion. The final token of each stimulus was selected such that the three final stimuli within a triplet were as acoustically similar as possible except for C_1_. The final stimuli were normalized to the same loudness and matched for peak intensity. The acoustic durations of the entire stimulus and the approximate duration from the stimulus onset to C_1_ onset judged by two phoneticians are as follow: /izo/ = 398 ms (stimulus duration), 134 ms (first syllable duration) ms., /ivo/ = 394 ms, 134 ms., /iᴣo/ = 394ms., 134ms., /ød^ɔ᷉^ t/= 667 ms., 131 ms., /øb^ɔ᷉^ t/ = 659 ms., 114 ms., /øg^ɔ᷉^ t/ = 657 ms., 114 ms.

### Procedure

Each participant was individually tested over two sessions separated by one week. The first session began with the novel word learning phase followed by the unattentive EEG oddball paradigm recording. Only the unattentive oddball task was repeated in the second session without additional learning.

### First session (Day 1)

#### Novel word learning phase

The learning phase was separated into four parts described below and lasted a total of 30 min.

#### Part 1

At the beginning of the learning phase, participants were simultaneously exposed to the spoken and written forms of the six novel words from the two triplets. The spoken inputs were presented through earphones while the written inputs were visually presented on a computer screen. The associations between the spoken and written forms corresponded to the description in Table 1. Each novel word was presented eight times, leading to a total of 48 stimuli presented in a random order.

#### Part 2

After the initial exposure, participants performed a dictation task to strengthen the learning of novel words’ spellings. During this task, the six novel words were orally presented one by one in a random order and the participants were required to write down each novel word using the spelling that they had learned. The list of six novel words was repeated three times.

#### Part 3

Following the dictation task, the association between sounds and spellings were further reinforced using a letter detection task. On each trial, a letter was presented on-screen for 750 ms and immediately after, one of the six novel words was presented aurally. Participants had to decide as fast and as accurately as possible whether the letter was present in the novel word’s spelling. Feedback (correct vs. incorrect response) and the correct spelling of the novel word were provided on the computer screen after each trial. The task comprised six blocks of 36 trials (six repetitions per novel word within each block). On half of the trials, the letter was present in novel word’s spelling.

#### Part 4

At the end of the learning phase, the dictation task was conducted again on the same six novel words to ensure that their spellings had been correctly acquired.

#### Unattentive oddball paradigm with EEG recording

The two triplets of novel words were presented in two separated blocks. Each block contains 960 repetitions of the standard stimulus and 120 repetitions of each of the deviant stimuli, corresponding to 11.1% of all trials. The stimuli were randomly presented with the constraint that there were at least three standard stimuli between two deviants. The inter-stimulus interval varied between 500 and 700 ms. Participants were informed that they would hear the novel words presented repeatedly via headphones. Throughout the auditory task the participants watched silent documentaries (without subtitles to avoid activation of orthographic representations) and were asked to ignore the auditory stimuli. EEG was recorded during the entire session and participants were instructed to remain as still as possible. The presentation order of the two blocks was counterbalanced across participants. The session lasted approximately 45 minutes.

At the end of the session, the participants performed the dictation task on the six novel words to ensure that their spelling knowledge remained accurate after the oddball paradigm phase.

### Second session (Day 8)

For each participant, the second session took place one week after the first one. The participants carried out the same unattentive oddball task with EEG recording. No additional novel word learning was performed. After the oddball paradigm, the dictation task was conducted.

For all tasks, stimulus presentation and behavioral data acquisition were controlled by E-prime 2.0 software (Psychology Software Tools, Pittsburgh, PA).

### Data recording and analysis

#### EEG recording

During the experiment, participants were seated in a soundproof and dimly lit experimental booth at 100 cm from a 13-inch laptop. This configuration allowed participants to visualize the full screen without inducing important eye movements. The EEG system (256-channel EGI Geodesic HydroCel) was continuously recording during the unattended oddball task at a sampling rate of 500Hz using the EGI software Net Station 5. This system uses the vertex (Cz) as the recording reference. The impedance of the electrodes was checked prior to recording and was kept below 40 kOhm. The two triplets of novel words were recorded in two separate runs and the order of presentation was counterbalanced across participants. After EEG acquisition, the electrode positions were localized by photogrammetry via EGI’s GPS sensor digitization system and their GPS 3.0 Solver.

#### Preprocessing of EEG data

Offline preprocessing was conducted using Brainstorm^59^ (neuroimage.usc.edu/brainstorm). Continuous data were bandpass filtered between 0.3 and 40Hz using a FIR Kaiser filter (order 7252) with a stopband attenuation set to 60dB. The EEG electrodes located on the face and on the neck (75 channels) were excluded from further analysis. For each participant. Power spectrum density was computed to control the overall quality of the recording and to identify bad channels. After bad channel rejection, the signals were re-referenced to the average of the mastoids. For one participant, one mastoid channel was identified as bad, therefore a neighboring channel was used for re-referencing.

The continuous data were visually inspected in order to identify segments of signal containing artifacts (e.g. muscle artifacts, movements, except eye blinks) and remove them from subsequent analysis. This visual inspection was performed on non-labelled signals. So, it was not biased by the events of interest. Eye blinks artifacts were first automatically detected and corrected using signal space projection (SSP). It should be noted that in order to create the SSP projector, eye blinks which co-occurred with events of interest or with other types of artifacts (+-200 ms) were removed before constructing the SSP projector.

Epochs from -200 to 800 ms relative to the onset of the first syllable were extracted and baseline corrected by DC-offset subtraction (-200 to 0 ms). Epochs containing any segment of artifacts were rejected. A total of 11.04% of the trials were excluded (deviant: 11.11%, standard: 11.03%).

#### ERP analysis

One critical issue for the present study was to ensure that any difference that might be observed between the MMNs generated by orthographically congruent and by orthographically incongruent deviants was due to novel words’ spellings rather than any acoustic or phonetic differences that might exist between the two deviant stimuli. To this end, the data obtained in the two groups of participants were combined in all subsequent analyses such that the spoken forms of both deviant stimuli were presented in both orthographically congruent and incongruent conditions (Table 1) and only the spelling form differed.

Since the first syllable of the standard stimulus from Triplet 2 was 17ms longer than the first syllable of the deviants, the signals of the deviants were shifted by 17 ms prior to computing the MMN.

Three subsets of electrodes were defined by selecting three clusters of 15 electrodes around FCz, Cz and Pz. The anterior-central cluster contains E036, E029, E022, E014, E005, E224, E030, E023, E015, E006, E215, E024, E016, E007, E207. The central cluster contains E008, E198, E185, E144, E131, E090, E080, E053, E044, E017, E009, E186, E132, E081, E045. The posterior-central cluster contains E100, E101, E129, E099, E110, E119, E128, E141, E109, E118, E127, E140, E117, E126, E139.

In the first stage, we functionally identified the peak of the MMN. For each triplet, the grand average signal was computed across all subjects, conditions (i.e. incongruent, congruent), sessions and electrodes clusters, separately for standard and deviant. The grand average MMN was obtained by subtracting the averaged signal of the standard stimuli from the averaged signals of the deviant stimuli. Then, for each triplet, the peak was detected on the grand average MMN over a theoretical time-window (200 to 400 ms, taking into account the acoustic duration of the first syllable).

In the second stage, at the individual level, the MMN peak for each triplet falling within a 100ms time-window centered around the group-level MMN peak was identified. This procedure was conducted separately for each condition, each session and each cluster. For each detected peak, three measures were extracted: 1) amplitude, 2) latency, 3) Area Under Curve (AUC) corresponding to 40 ms around the peak. The same procedure was conducted at both group and individual levels within the baseline period, relative to the first syllable onset, from -200 to 0 ms; this allowed us to compare the presence of the MMN against the real baseline signal rather than against zero.

In order to combine the data from the two triplets in the same analysis, we applied a temporal shift of the T0 to realign the signals to the onset of the second syllable (this resulted in a 134 ms and 114 ms shift for the Triplet 1 and Triplet 2, respectively). Hence, the MMN latency reported here is relative to the onset of the second syllable.

Statistical analyses were conducted in two stages. First, we verified the existence of the MMN within each of the three electrode clusters. To this end, we conducted t-tests comparing the amplitude of the averaged MMN (all triplets, conditions and sessions combined) extracted from the time-window of interest to the signal extracted from the baseline period. The procedure was applied separately on the signal obtained from the three electrode clusters. Only those clusters presenting a significant MMN were retained for the subsequent analyses. Second, we examined the effects of orthographic congruency and experimental session, using R software (R Core Team, 2014), with a linear mixed-effects model (LME). The model was fitted with the lme4 package (Bates et al., 2014) and the p values (Type III: marginal sum of squares) were computed with the lmerTest package (Kuznetsova et al., 2013). The model was applied to peak amplitude, peak latency and AUC. Orthographic congruency, session, cluster and their interactions were considered as fixed factors. Participants, triplets and their interactions were considered as random intercepts.

#### Source localization analysis

To identify the cortical origin of the orthographic congruency effect observed at the scalp electrodes, an exploratory source reconstruction analysis was conducted using Brainstorm software^59^. To this end, the coordinates of the 256 electrodes and three landmarks (LPA, RPA and Nasion) were extracted for each participant using the GPS 3.0 Solver. For each participant the ICBM152 MNI template anatomy was warped such that the scalp matched the head shape defined by the digitized head points. To estimate the forward (head) model, the boundary element method (BEM) from OpenMEEG was applied within Brainstorm with no constraint on the orientation of the sources. Noise covariance matrices were calculated on a pre-stimulus baseline (i.e., -300 to 0 ms) of all trials from all conditions for each recording run (two per participant). The dynamic statistical parametric mapping (dSPM) method was used to estimate the neural activity at the cortical level with 15,002 vertices both for the averages of standard and deviant stimuli and the results obtained on the standard stimuli were subtracted from those of the deviant stimuli from the congruent and incongruent condition, respectively. In order to identify the cortical source of the orthographic congruency effect observed at the sensor level, the peak latency of the MMN detected at the sensor level was used for the source level analysis. Hence, within the time-window of interest, a source reconstruction was obtained for each triplet, subject, condition, and session.

Since the source model was computed with unconstrained orientation, each vertex presented three time series (x,y,z,). To reduce the three dimensions to one single dimension, a PCA was computed for each vertex. The absolute value was then output to SPM12 (https://www.fil.ion.ucl.ac.uk/spm/) for further statistical comparisons. In SPM12, a paired t-test examining the effect of orthographic congruency observed at the sensor level was applied to source activity maps with cluster and triplet as covariates.

## Acknowledgements

This work was supported by the French Ministry of Research: ANR-13-JSH2-0002 and ANR-19-CE28-0001-01 (to C.P.), ANR-16-CONV-0002 (ILCB), ANR-11-LABX-0036 (BLRI) and the Excellence Initiative of Aix-Marseille University (A*MIDEX). It was performed in the Centre d’Expérimentation de la Parole (CEP), Laboratoire Parole et Langage. We warmly thank Dr. Agnès Trébuchon for taking medical responsibility during the study. Center de Calcul Intensif d’Aix-Marseille is acknowledged for granting access to its high-performance computing resources.

## Data availability

The datasets generated and/or analyzed during the current study are not publicly available since the ethical approval for this study does not include permission to share data in a public data repository but they are available from the corresponding author on reasonable request.

## Author contributions

C.P. A-S.D., and D.B. designed research, collected data; A-S.D and S.W. analyzed and visualized data; C.P., A-S.D. and S.W. wrote the original manuscript, All authors reviewed the manuscript.

