## supplementary material for "Learning to read transforms phonological into phonographic representations: Evidence from a Mismatch Negativity study"

**Table 1S.** Brain regions showing the orthographic congruency effect in the second experimental session. Hemisphere, MNI coordinates (X, Y, Z), T statistics for the peak and cluster size (number of vertices) are reported. Brain regions were identified using Automated Anatomical Labeling atlas 3 (AAL3)[1].

| Brain region | Hemisphere | X | Y | Z | T | Number of vertices |
|---|---|---|---|---|---|---|
| *Regions showing higher activity in the incongruent condition* | | | | | | |
| Middle Frontal | L | -23 | 20 | 47 | 4.27 | 28 |
| Superior Parietal | L | -34 | -51 | 72 | 5.44 | 21 |
| Middle Temporal | L | -64 | -37 | 4 | 4.74 | 14 |
| | L | -62 | -31 | -2 | 4.54 | 13 |
| *Region showing lower activity in the incongruent condition* | | | | | | |
| Fusiform | L | -53 | -59 | -24 | 4.17 | 17 |
| | L | -47 | -58 | -23 | 4.26 | 12 |
